# LiFT: Live foci tracking for quantitative analysis of DNA damage dynamics

**DOI:** 10.64898/2026.07.17.739177

**Authors:** Tijmen H. de Wolf, Pleun A.M. Engbers, Justine Perrin, Krijn H. van der Steen, Sam F.B. van Beuningen, Ihor Smal, Julie Nonnekens

## Abstract

Quantitative analysis of radiation induced DNA double strand breaks (DSBs) and their repair is essential for understanding and eventually contributing to improving radiation-based cancer therapies. Using live-cell microscopy, the formation and resolution of DSBs over time can be followed in individual cells through tracking of foci formed by accumulation of DSB repair proteins. However, manual analysis of such time-lapse datasets is a tedious time-consuming task that is prone to operator bias, affecting the reproducibility. Here, we present LiFT, an automated image analysis pipeline, specifically designed for robust quantification of DSB kinetics in live-cell imaging experiments. To quantify DSB kinetics, our pipeline first segments and tracks cell nuclei without requiring a nuclear stain. After correcting for inter-frame motion through image registration, automatic detection and tracking of foci within these nuclei enables direct quantification of the dynamics of individual repair events. Multiple algorithmic options were implemented for each step of the pipeline, ensuring more general applicability to potentially different imaging setups and applications. We evaluated the pipeline using PLC/PRF/5 cells and demonstrated its generalizability on U2OS-SSTR2 cells. Our results show that LiFT enables reproducible and scalable quantification of DSB dynamics, providing a broadly applicable framework to analyse live-cell imaging data in cancer research. To improve the adoption of LiFT, we made it available as an open-source Python package and provided a graphical user interface to select different methods and adjust method related parameters.

## Introduction

Radiotherapy is a cornerstone therapy for cancer as curative or palliative treatment option^1^. Its ionizing radiation damages the DNA and of the different types of induced DNA damage, double strand breaks (DSBs) are the most detrimental for cellular survival and lead to cell death if left unrepaired^2^. Cells possess an extensive network of DNA repair pathways, called the DNA damage response (DDR), to repair DNA damage induced by ionizing radiation^3^. The ability of cells to successfully repair the damage ultimately controls their fate: persistent or irreparable damage can result in programmed cell death or senescence, whereas efficient repairs permit survival and proliferation^2,4^. After DSB induction, the DDR initiates a cascade of responses that recognize the lesions and enable the spatial organization of repair factors. Among the earliest events is the phosphorylation of histone H2AX at the sites flanking the DSBs, followed by the recruitment and accumulation of p53-binding protein 1 (53BP1), after which pathway specific proteins will be recruited to finalize the repair^5^. The spatial accumulation of these proteins results in discrete subnuclear structures with high local concentration. Immunofluorescence staining or tagging of these proteins therefore allows the proteins to be visualised using fluorescence microscopy as so-called foci^2,5,6^.

Quantification of the DNA damage foci in confocal microscopy images is widely regarded as the gold-standard method for studying radiation biodosimetry and the DDR^7–10^. While fluorescence imaging of fixed cells provides a static snapshot at predefined time points, live-cell microscopy allows individual cells to be followed over time, capturing the full temporal dynamics of foci formation and resolution, providing deeper insights into the kinetics of the DDR^11–13^. This is particularly important for therapies with protracted induction of DNA damage (such as chemotherapy or targeted radionuclide therapy, a form of internal radiotherapy) where induction and repair of DSBs occurs simultaneously. Using adequate time-sampling between frames, individual foci can even be followed over time^11–14^, providing a biomarker for the repair time of the individual DSB^13^.

Despite these distinct advantages offered by live-cell imaging for studying DNA repair kinetics, it also introduces several technical challenges. In contrast to the imaging of fixed-cells, where a field of view (FOV) is acquired only once, live-cell imaging repeatedly images the same cells over extended periods. This repeated exposure to the excitation light can result in photobleaching of the fluorescent proteins and subsequent loss of the foci signal^11,12,15^. Moreover, depending on the wavelength of the excitation light source, the illumination itself may induce cellular damage, known as phototoxicity^15^. To mitigate these adverse effects, laser power is typically kept low, which reduces the signal-to-noise ratio of the acquired images^16,17^. Reducing the acquisition frame rate is an alternative strategy to limit exposure, but this comes at the cost of temporal resolution. Indeed, due to movement of the cells and their foci over time, longer intervals between frames can substantially reduce the tracking accuracy and in extreme cases render tracking infeasible^12,18,19^. Consequently, live-cell imaging data are often characterized by low signal-to-noise ratios, complicating the interpretation and analysis^17,20^.

Beyond imaging challenges, live-cell microscopy generates large datasets that are difficult to analyse manually. Such data are still frequently quantified manually, which is tedious, extremely time-consuming, and prone to inter-observer variability^8,10,11,21,22^. Live-cell imaging further amplifies these issues due to the large number of time points and need for precise positional information for downstream analyses such as tracking of foci. Consequently, manual quantification of DSB repair kinetics presents an prohibitive annotation burden, underscoring the need for automated analysis tools^10^.

Despite this need, dedicated image-processing pipelines for extracting DNA repair kinetics from live-cell microscopy experiments remain scarce. As a result, many studies still rely on manual analysis, including hand-tracking of individual foci^13,23^, often yielding only limited number of trajectories (e.g., 200 in^13^ and 188 in^23^ across all conditions). Other works performed live-cell imaging, but restricted the analysis to counting of the foci without tracking^11^. Although some studies report semi-automated approaches^24^ or automated pipelines^12,25^, these tools are either not publicly available or do not leverage recent advances in deep-learning based image analysis.

Here, we present a flexible and modular pipeline for Live Foci Tracking (LiFT) to analyse live-cell imaging data of DNA damage foci. LiFT integrates state-of-the-art deep-learning models to ensure robust performance across diverse imaging conditions. To enable analysis on a level of a single cell, we detect and track nuclei over time without requiring additional nuclear stains, which can interfere with DNA and the DDR^26,27^. Subsequently, we apply motion correction to stabilise the nuclei before detecting and tracking individual foci. Each stage offers multiple algorithmic options, and an interactive web-browser-based graphical user interface allows the user to select the desired methods and adjust the corresponding hyperparameters to optimally deal with the problems associated with their data (Supplementary Figure 1). The modular design also enables researchers to replace any component with custom or pre-trained models tailored to their specific imaging setup. By providing an accessible, adaptable and reproducible pipeline, we aim to facilitate broader use of live-cell imaging for studying DSB repair kinetics and to streamline data quantification.

## Methods: overview of the LiFT pipeline

A schematic overview of the LiFT pipeline is provided in Figure 1, a more detailed overview, including the available algorithms, is provided in Supplementary Figure 2. LiFT consists of two main stages: the detection and tracking of cell nuclei, followed by the detection and tracking of individual foci within each nucleus to characterize DSB induction and repair kinetics. The first stage comprises segmentation (step 1) and tracking (step 2), yielding single-cell trajectories over time. The tracked nuclei are then isolated from the FOV and aligned across frames by means of image registration (step 3) to compensate for cellular motion, producing “stabilized” single-nucleus image stacks. Using the stabilized stacks, foci are detected (step 4) and tracked (step 5), enabling the temporal dynamics of individual repair events to be quantified. To accommodate a broader range of imaging setups and experimental conditions, multiple algorithmic options are provided at each step.

**Figure 1.**
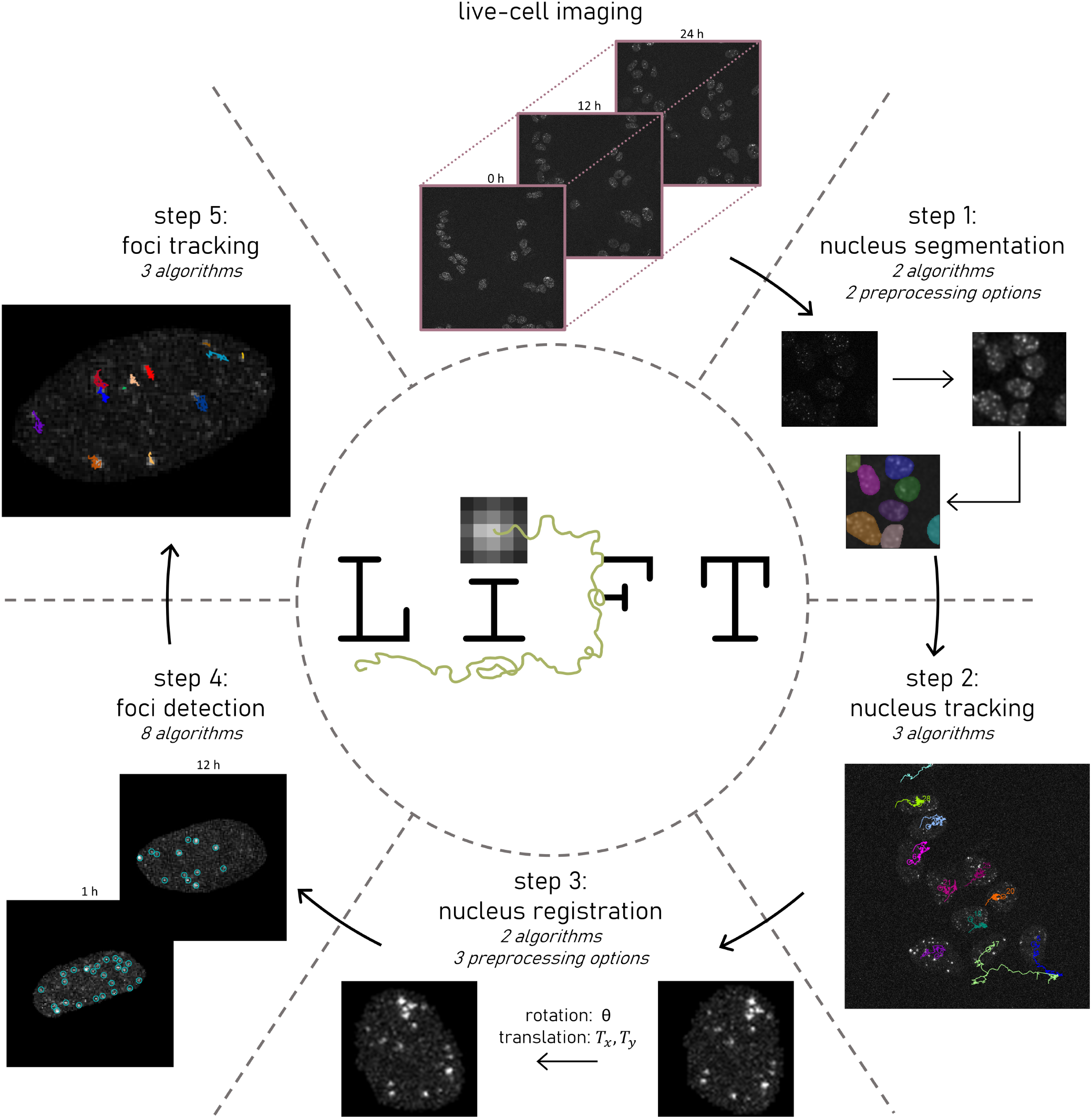
Schematic overview of the LiFT pipeline. Following data acquisition, nucleus segmentation and tracking are performed. The tracked nuclei are then isolated and image registration is applied to correct for motion between the frames. Subsequently, individual foci are detected and tracked. Multiple algorithmic options are available at each step, allowing users to select the algorithm best suited for their data.

To assess the generalizability of LiFT across cell lines and experimental conditions, we performed live-cell imaging for 24 hours on two distinct cell lines (PLC/PRF/5 and U2OS-SSTR2), each expressing an mClover endogenous fluorescently tagged 53BP1 protein. The PLC/PRF/5 cells were previously generated and validated by Perrin *et al*^28^ and the U2OS-SSTR2 cells were generated and validated for this study using the same protocol. The two cell lines were imaged under different treatment conditions (applied radiation doses of 0, 1.5 or 2.5 Gy for PLC/PRF/5 and 0 or 2 Gy for U2OS-SSTR2) and with different microscope acquisition parameters (imaging interval, laser power, detector gain and detector bandwidth), yielding data with distinct biological and imaging characteristics. Full details of the cell lines, irradiation, and imaging acquisition are provided in the Supplementary material (section 1.1).

### Step 1: Nucleus segmentation methods

The first step of LiFT is nucleus segmentation. Nuclear stains were not used because of their potential interference with the DNA and therefore with the kinetics of DSB repair^26,27^. For segmentation, LiFT uses Cellpose-V3^29^ and Cellpose-SAM^30^, the latter incorporating the Segment Anything Model (SAM)^31^. Since both methods were trained on images containing nuclear stains, we developed two preprocessing approaches that transform the available signal of the foci into a representation resembling a nuclear stain, aiming to improve segmentation performance. The first, termed “contrast” applies Gaussian smoothing followed by intensity clipping based on the image average intensity. The second, “wavelets” employs the isotropic undecimated wavelet transform (IUWT) with a B3-spline kernel to decompose the image into components of different sizes^32,33^. By removing small objects and high-frequency features, this suppresses the foci while preserving the background unbound 53BP1-mClover signal. To quantify nucleus segmentation performance, we manually segmented all PLC/PRF/5 nuclei in every frame of one FOV irradiated with 2.5 Gy of X-rays, yielding a total of 7288 manual masks.

### Step 2: Nucleus tracking methods

Following segmentation, the segmented masks are tracked to obtain single-cell trajectories over time. LiFT implements three alternative tracking algorithms. The first one is an intersection-over-union (IOU)-based tracker, which links nuclei across frames by computing the optimal global assignment that maximizes mask overlap. The second one is a greedy Nearest Neighbour Diffusion (NND) tracker, available through the SOS tracking plugin^34^. The third one is Trackastra, a state-of-the-art deep learning tracker that ranks among the top performers on the Cell Tracking Challenge linking benchmark^35^. We used the model provided with Trackastra, which was trained on the Cell Tracking Challenge dataset^36^. To quantitatively assess tracking performance, we manually tracked the ground-truth segmentations of the same PLC/PRF/5 FOV irradiated with 2.5 Gy of X-rays, yielding 37 trajectories with an average duration of 197 frames. Performance was quantified using the linking benchmark metrics defined by the Cell Tracking Challenge^36^.

### Step 3: Registration methods

Using the tracked nuclei, LiFT generates single-cell movies by cropping each nucleus from the full FOV across time. However, due to rotation and translation of the nuclei, direct tracking of foci is not reliable without motion correction. Similar to Jakob *et al* and Roobol *et al*, we applied rigid-body image registration to compensate for this motion^12,13^. LiFT includes two registration methods: StackReg^37^ and `elastix` ^38^, the latter is accessible through a Python wrapper we developed. StackReg only implemented the mean-square-error (MSE) as optimization metric, for `elastix` LiFT supports the MSE, mutual information (MI) and normalised cross-correlation (NCC) as optimization costs. Since live-cell imaging often results in noisy images that can degrade intensity-based registration, LiFT provides three optional preprocessing methods designed to enhance registration robustness by suppressing background signal and emphasizing the foci. The first is based on the IUWT, in which only objects within the expected foci size range are retained in the reconstruction. The second applies a Difference of Gaussian (DoG) filter, also tuned to the expected foci size. The third performs simple thresholding, setting signal below a certain threshold to zero while retaining the brighter 53BP1-mClover foci. These preprocessing steps ensure that registration is driven primarily by the foci, whose accurate alignment is essential for downstream tracking. In the StackReg implementation, rotation and translation are optimized exclusively on the preprocessed image and subsequently applied to the unmodified image. In contrast, `elastix` determines the transformation by jointly optimizing on both the preprocessed and original channels. Registration performance was quantified on the manually tracked nuclei, comprising 7251 frame pairs. We used both image-based metrics (the MSE and structural similarity index measure (SSIM)) and foci-based metrics, including the Chamfer distance^39^ and the median distance between detected foci. For the foci-based metrics, foci were detected using a DoG-based spot-detection algorithm.

### Step 4: Foci detection methods

To characterize the temporal behaviour of foci in live-cell imaging data, LiFT provides eight alternative detection algorithms. Most follow a standard two-step approach^20^, first enhancing the foci signal and subsequently identifying local maxima with a user-defined threshold. The following algorithms are included, with their associated parameters:

- *IUWT*: This method decomposes the image into components of different sizes^32,33^. The image is separated into *K ∈* `ℕ` wavelets scales, where the first scale predominantly captures noise. Each scale is thresholded by a shrinkage factor *f*_*W*_ using Jeffreys’ non-informative prior, to suppress noise while retaining the foci^40^. User-controlled parameters are the number of scales *K*, the shrinkage factor *f*_*W*_ and which scales to retain/include in the reconstruction.
- *Laplacian of Gaussian (LoG)*: This filter removes local background and noise while preserving spot-like structures such as foci^20,41^. It takes a single user-defined parameter *σ* (the standard deviation of a Gaussian), which should match the size of the foci.
- *Hessian*: This method uses the determinant of the Hessian matrix to estimate local image curvature, enabling discrimination of high-curvature foci from noise and background. Prior to curvature estimation, the image is smoothed with a Gaussian with *σ*, which is the sole user-controlled parameter.
- *Top-Hat*: The Top-Hat filter, commonly used in foci detection literature^42,43^, removes local background by applying grayscale opening with a disk-shaped structure element of radius *R*. The result of this opening is subtracted from the original image, yielding an image in which structures smaller than the structuring element are preserved. Prior to applying the Top-Hat operation, the image is smoothed using a Gaussian filter with *σ* .
- *H-dome*: The H-dome transform enhances bright local maxima by suppressing smoothly varying background without relying on a size criterion^20,44^. It extracts local maxima that rise at least a user defined minimum intensity *h* above their surroundings. The method performs grayscale morphological reconstruction on an image from which *h* has been subtracted, using the original image as the reconstruction mask. The H-dome image is then obtained by subtracting the reconstructed result from the original image. the value *h* is the only user-defined parameter for this method.
- *H-dome + LoG*: This method is based on the detection scheme introduced by Smal *et al*^45^, where the H-dome transform is applied to a LoG filtered image. The LoG filter, parameterized by *σ*, removes local background and enhances the signal of the foci. The output of the H-dome transformation is raised to the power *s*, to produce a more narrowly peaked signal. This method therefore has three user-controlled parameters, *h, σ* and *s*.
- *Maximum Possible Height Dome (MPHD)*: This method extends the H-dome approach by computing an adaptive *h* value for each candidate spot, making it better suited for images in which no single global value of *h* is appropriate^46^. First, noise is suppressed by Gaussian smoothing with a filter of width *σ* . An initial H-dome transform with height *h*_*init*_ identifies candidate local maxima. For each maximum, the algorithm determines the maximum possible dome height by searching within a radius *R* for the minimum surrounding intensity. These locally optimized heights define the reconstruction mask used to compute the final MPHD-enhanced image.
- *Spotiflow*: Finally, Spotiflow is a state-of-the-art deep-learning spot detector^47^. Spotiflow was trained on a broad range of imaging conditions, yielding a generally applicable model that performs well in noisy conditions and on previously unseen datasets or data from different domains. The network predicts a vector field pointing to the centre of the nearest spot and an associated heatmap indicating the probability of a spot at each location. Unlike the other methods presented above, Spotiflow has no further user-defined parameters.

### Step 5: Tracking methods for foci

To quantify the dynamics of foci formation and resolution, LiFT performs automatic tracking of the detected foci and provides three alternative tracking algorithms. The first is a Global Nearest Neighbour (GNN) tracker, which links detections across frames by solving a global assignment that minimizes the total distance between detections, following the formulation introduced in Jaqaman *et al*^48^. The second is the Non-iterative Greedy Multi-frame Assignment (NGMA) tracker, available through the SOS tracking plugin^34^. Like the GNN tracker, it links detections based on spatial proximity, but incorporates a multi-frame buffer that enables it to consider associations several frames ahead. The third is Trackastra^35^, the same deep-learning-based tracker used for nucleus tracking; for foci, the “general” pre-trained model was selected, which was trained on a broad range of datasets including those from the particle tracking challenge^49^. Tracking performance was evaluated using the metrics defined in the particle tracking challenge^49^. To ensure methods were not underperforming due to poor parameter choices, we performed a joint parameter sweep across the foci-detection and tracking algorithms and selected the parameters yielding the highest Jaccard similarity coefficient (JSC) for the resulting tracks. For conciseness, the performance was evaluated using three representative detectors (even though LiFT supports additional options as well): i) the LoG detector, widely used in tools such as TrackMate^50^, ii) the Top-Hat detector, frequently used in the foci detection literature^42,43^, and iii) Spotiflow, a state-of-the-art deep-learning spot detector^47^. Automatic tracking results were compared to a manual ground truth derived from two of the manually tracked nuclei (registered using StackReg without a processing function), in which all foci were tracked across all 288 frames (24 hours), yielding 294 ground-truth trajectories and 7710 annotated detections.

### Statistical analysis

Once foci have been detected and tracked, LiFT enables direct comparison of repair kinetics across experimental conditions, such as different irradiation doses or modalities. To support this, the pipeline includes built-in statistical testing for differences in track-derived metrics: differences in mean dwell times are assessed using a non-parametric permutation test, and confidence intervals (CIs) are obtained by bootstrap resampling with 5000 resamples using the percentile method. Throughout this study, the statistical significance was defined as *p* < 0.05.

### Availability of LiFT

To facilitate adoption by the community, LiFT is distributed as an open-source Python package, freely available at https://github.com/UU-cellbiology/LiveFoci and PyPI https://pypi.org/project/live-foci/. Multiple interfaces are provided to accommodate users with different levels of programming experience: a browser-based graphical user interface (Supplementary Figure 1), a Jupyter notebook exposing the individual processing steps, a PyPI package for integration into custom pipelines and a command-line interface for scripted execution. To support reproducibility across different computing platforms and operating systems, a modular Docker image is provided on Docker Hub https://hub.docker.com/r/krijns/lift. Finally, to ensure transparency and reproducibility of the analysis, parameter files documenting the settings used at each step are automatically saved alongside the output, allowing other researchers to reproduce the tracking results from the same input data.

## Results

### Nucleus segmentation

The first stage of our pipeline is the segmentation of nuclei based solely on the presence of foci, as nuclear stains can alter repair kinetics^26,27^. We incorporated two state-of-the-art segmentation models: Cellpose-V3^29^ and Cellpose-SAM^30^, the latter employing a substantially larger model and therefore impractical for CPU-only execution. When applied directly to raw foci images, Cellpose-V3 resulted in poor segmentation performance (Table 1 and Figure 2A). To enable accurate segmentation even in the absence of a GPU, we developed two preprocessing routines that transform the signal of the foci into an image resembling a nuclear stain (Figure 2B, C, E, F). The proposed contrast and wavelet filtering improved Cellpose-V3 performance, increasing precision from 2.6% on the raw data to 80.1% and 78.9% respectively. Cellpose-SAM was also evaluated in combination with the preprocessing algorithms. However, it already achieved a precision of 93.5% and a Jaccard index of 0.838 without the preprocessing step (Figure 2D). Cellpose-SAM substantially outperformed Cellpose-V3 on both metrics (Table 1).

**Table 1.**
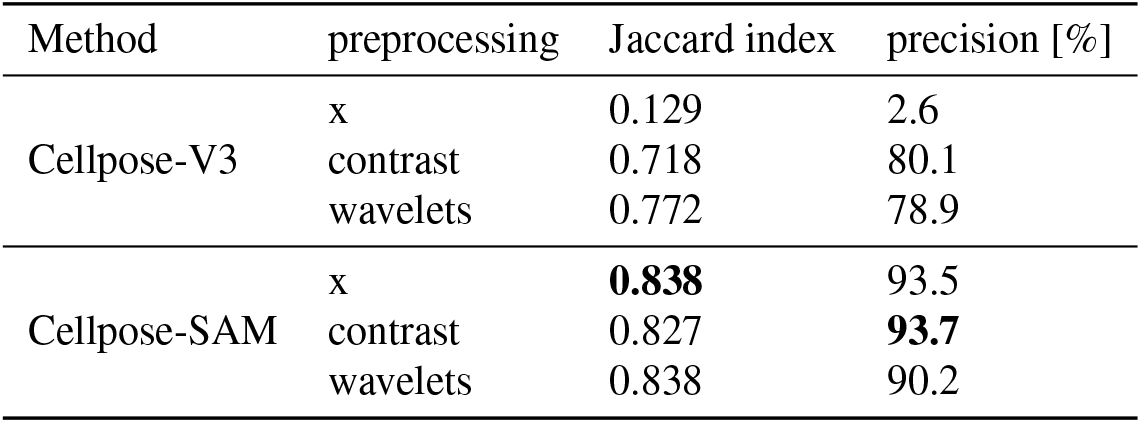
Quantitative evaluation of the cell segmentation performance for Cellpose-V3 and Cellpose-SAM in combination with different preprocessing functions. The “x” indicates no processing function was applied. Numbers in bold indicate the best performing method for each metric.

**Figure 2.**
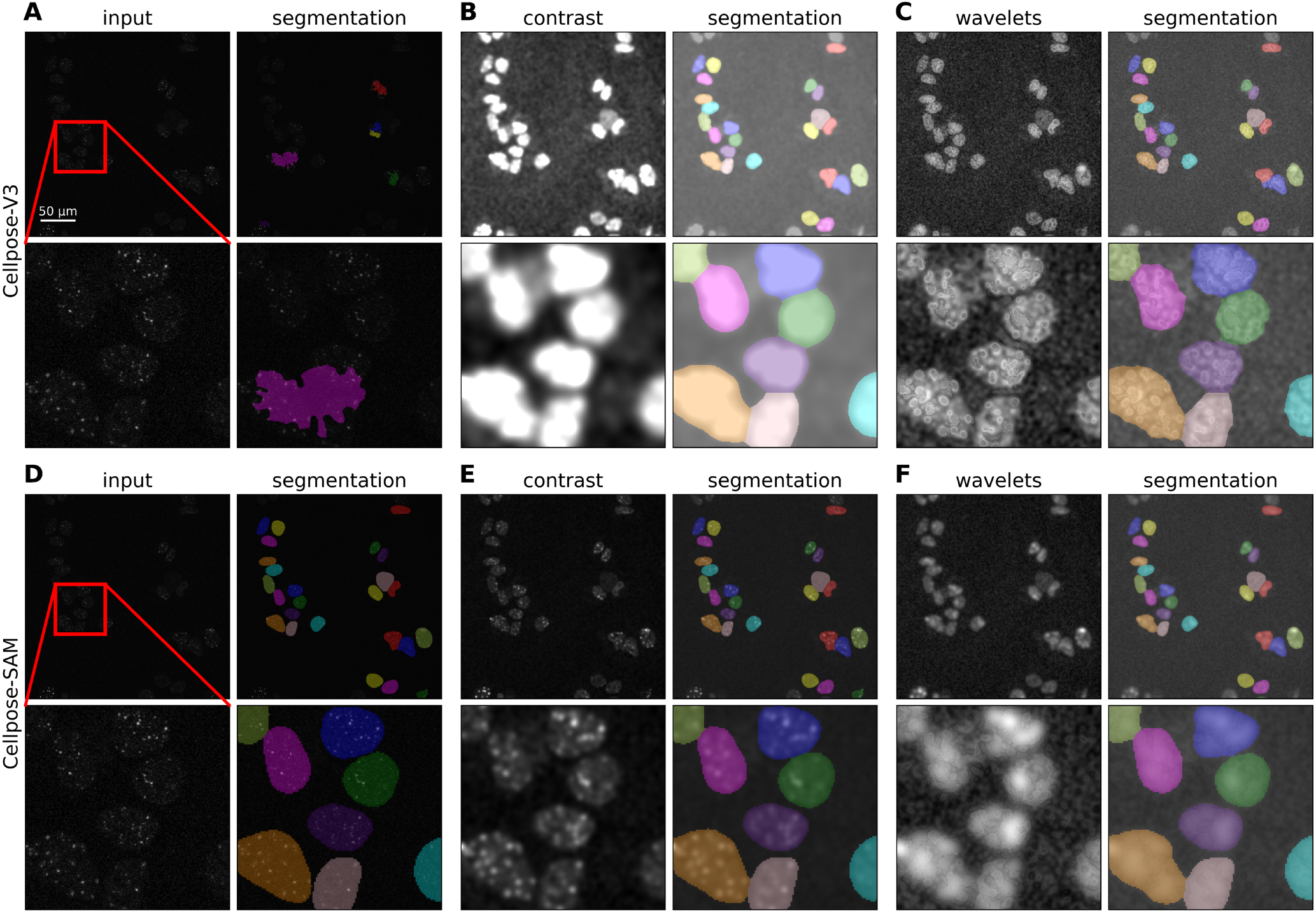
Representative images of the comparison of the available segmentation algorithms (step 1). Each panel shows the full field of view and a zoom in of the region indicated in red in the first image. (**A, D**) Raw image and the segmentation output of Cellpose-V3 (A) and Cellpose-SAM (D). (**B, E**) Result after contrast adjustment the input image and the segmentation result of Cellpose-V3 (B) and Cellpose-SAM (E) on the contrast adjusted image. (**C, F**) Resulting image after wavelet filtering the input image and segmentation using Cellpose-V3 (C) and Cellpose-SAM (F).

### Nucleus tracking

Using the segmented nuclei masks, we generated single-cell trajectories employing the different tracking algorithms implemented in LiFT: the IOU tracker, the nearest-neighbour^34^ tracker and Trackastra^35^. To evaluate performance under different segmentation conditions, we tracked nuclei using masks generated by Cellpose-V3 with contrast preprocessing (Figure 3A), representing CPU-limited scenarios, and masks produced by Cellpose-SAM (Figure 3B), which achieved the highest Jaccard index on this dataset. The NND tracker achieved the best overall performance on the Cellpose-SAM segmentations, outperforming the other trackers on three out of the four considered metrics (Table 2). However, compared to the other tracking algorithms the differences were minimal. To disentangle the tracking errors from the segmentation errors, we repeated the experiment using the ground-truth masks as input to the tracking algorithms. All three algorithms achieved a score of 1.0 on all metrics (TRA, CT, TF, LNK), indicating that tracking errors in this dataset predominantly stem from imperfect nucleus segmentation rather than from the tracking algorithms themselves.

**Table 2.** Quantitative evaluation of the cell tracking performance based on the segmentation masks from Cellpose-V3 with contrast adjustment as preprocessing step or the segmentations masks of Cellpose-SAM. Numbers in bold indicate the best performing method for each metric.

| Segmentation | Tracking method | TRA | CT | TF | LNK |
| --- | --- | --- | --- | --- | --- |
| Cellpose-V3 | IOU tracker | 0.908 | 0.078 | 0.750 | 0.818 |
|  | NND | 0.904 | 0.076 | 0.750 | 0.815 |
|  | Trackastra | 0.909 | 0.076 | 0.750 | 0.819 |
| Cellpose-SAM | IOU tracker | 0.974 | **0.247** | 0.917 | 0.965 |
|  | NND | **0.975** | 0.244 | **0.925** | **0.966** |
|  | Trackastra | **0.975** | 0.217 | 0.916 | **0.966** |

**Figure 3.**
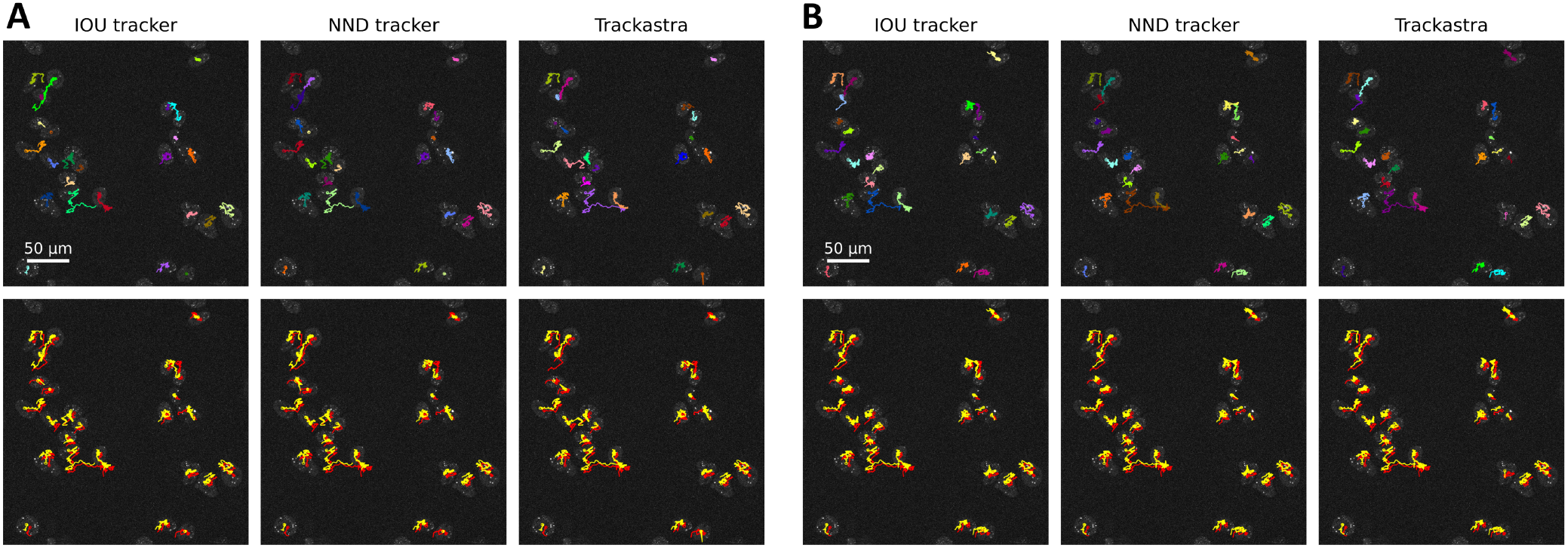
Performance of the three available cell tracking algorithms (step 2). (**A, B**) Tracking results on the segmentation masks of Cellpose-V3 (A) and Cellpose-SAM (B). The Top panel indicates the tracks, the bottom panel shows the comparisons of the tracking result (yellow) to the ground truth tracks (red). In the bottom panel, the ground-truth and predicted tracks were displayed with a small spatial offset to prevent visual overlap. For displaying purposes, the contrast of the images was adjusted in this figure.

### Nucleus registration

Based on the single-cell trajectories, we generated time-lapse sequences for each individual nucleus by isolating their spatial regions from the full FOV. To correct for displacement of nuclei between frames, we have applied motion correction using image registration. For this purpose, LiFT includes both StackReg^37^ and `elastix` ^38^. Since these methods operate on all pixels and our primary goal is to align the foci, we evaluated three processing functions to suppress background signal and enhance the foci prior to registration. To evaluate registration performance independently of segmentation or tracking errors, we used the ground-truth nuclei masks and trajectories. StackReg achieved the best alignment of the foci, as reflected by the distance-based metrics (Table 3, Figure 4), and required only 2.5 seconds to register a stack with 288 frames of 106 by 106 pixels compared to 5 minutes for `elastix` on an AMD Ryzen 9 5950X CPU. Conversely, `elastix` achieved slightly but non-significantly superior image-based registration metrics.

**Table 3.**
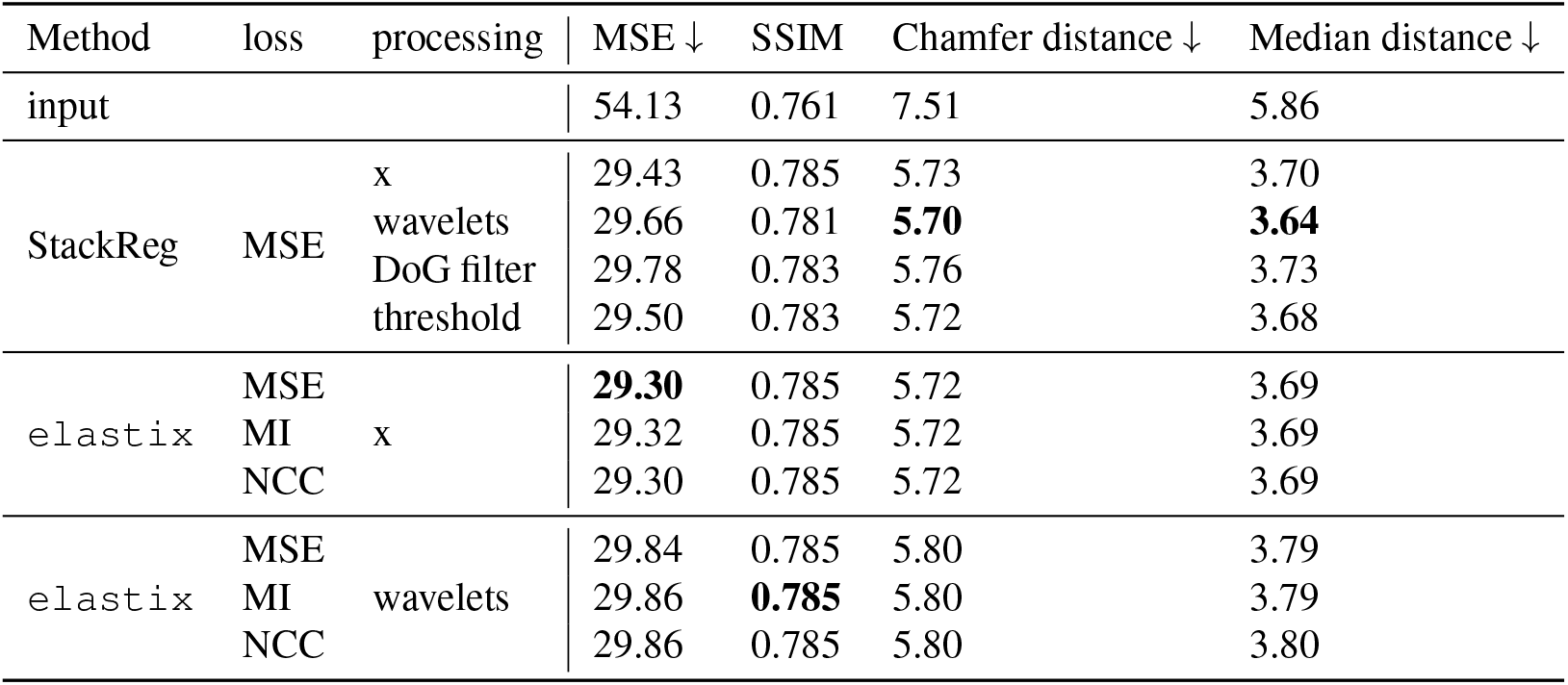
Quantitative evaluation of the registration methods. The “x” indicates no processing function was applied. Numbers in bold indicate the best performing method for each metric. All scores used to quantify the performance of the different algorithms should be maximized unless specified otherwise with a downward arrow: *↓*.

**Figure 4.**
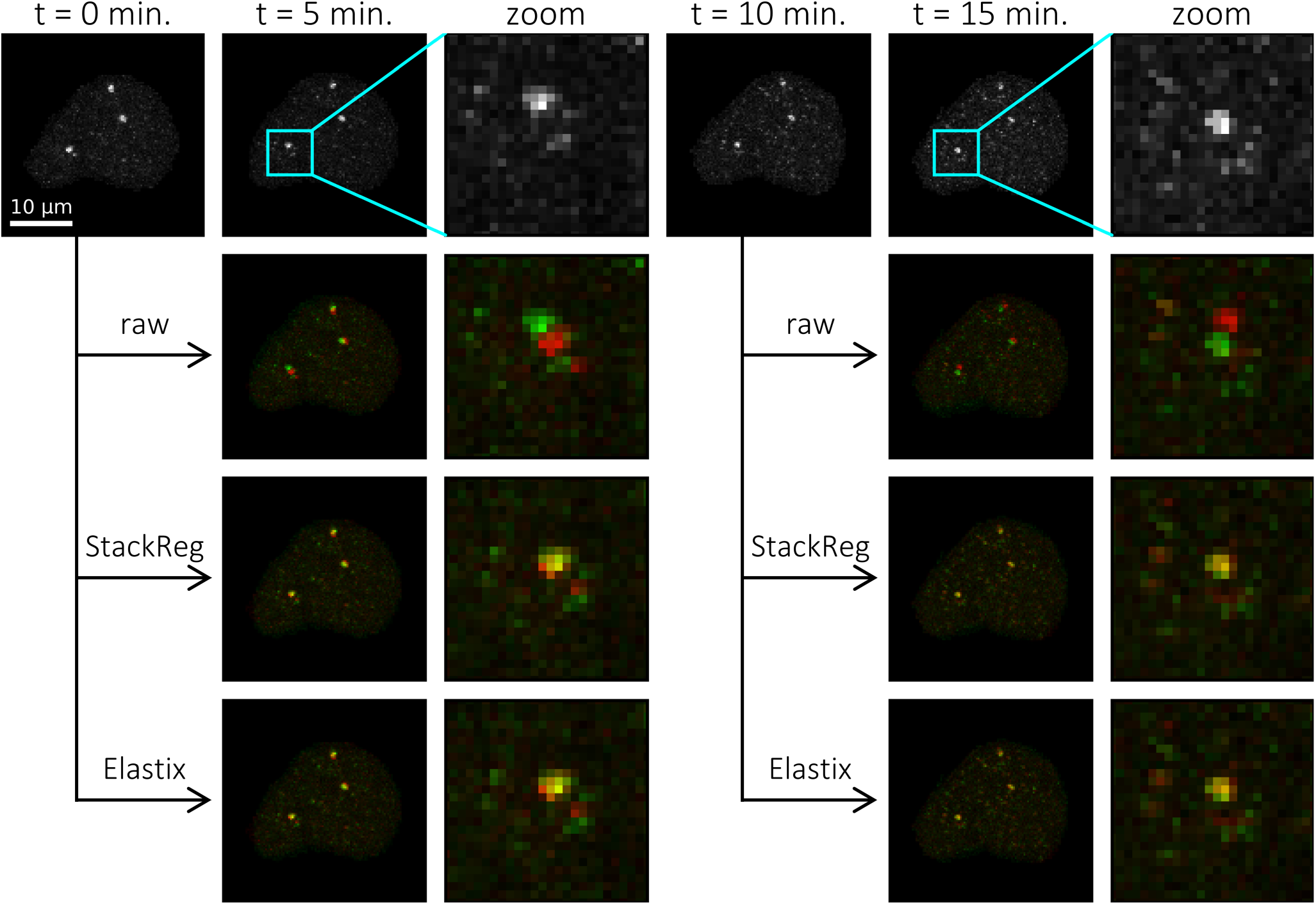
Visual representation of the registration (step 3) results for StackReg and `elastix` with MSE optimization. The top row depicts the input images. The rows below it show an overlay of the previous frame (red) and the current frame (green) for the input (row 2) and the result after registration with StackReg (row 3) and `elastix` (row 4). Column 3 and 6 contains a zoom-in of the indicated square region (cyan) from the previous column.

### Detection of foci

Detection of foci in LiFT is evaluated using eight available detectors. We jointly optimized detector and tracker combinations instead of assessing them in isolation and evaluated the performance of three representative detectors. The LoG detector underperformed substantially relative to the other detectors obtaining a maximum JSC for the detections of 0.347. We found the Top-Hat achieved the best detection score with a JSC of 0.557, closely followed by Spotiflow reaching a score of 0.518 (Figure 5A, Table 4).

**Table 4.**
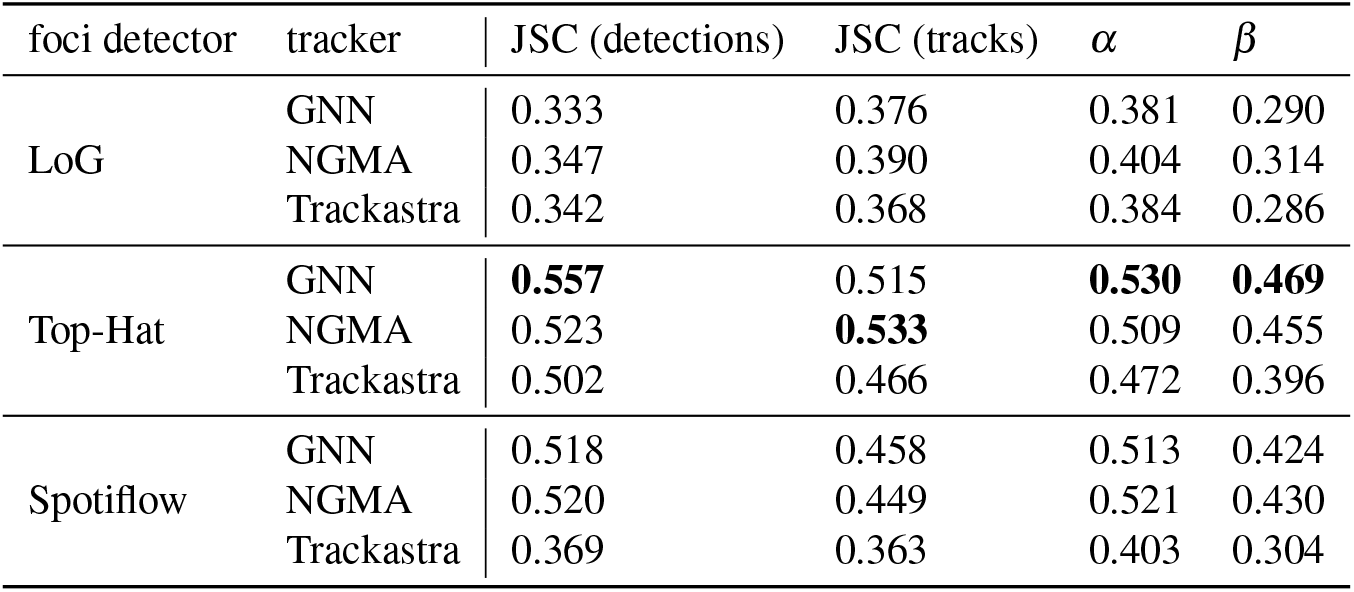
Quantitative evaluation of the detection and tracking performance of foci. Three representative detectors were selected, and all possible detector–tracker combinations were evaluated. Optimal parameters for each detector and tracker were determined via a parameter sweep, selecting the configuration that maximized the Jaccard similarity coefficient (JSC) of the resulting tracks. The values in bold indicate the best-performing method for each metric.

**Figure 5.**
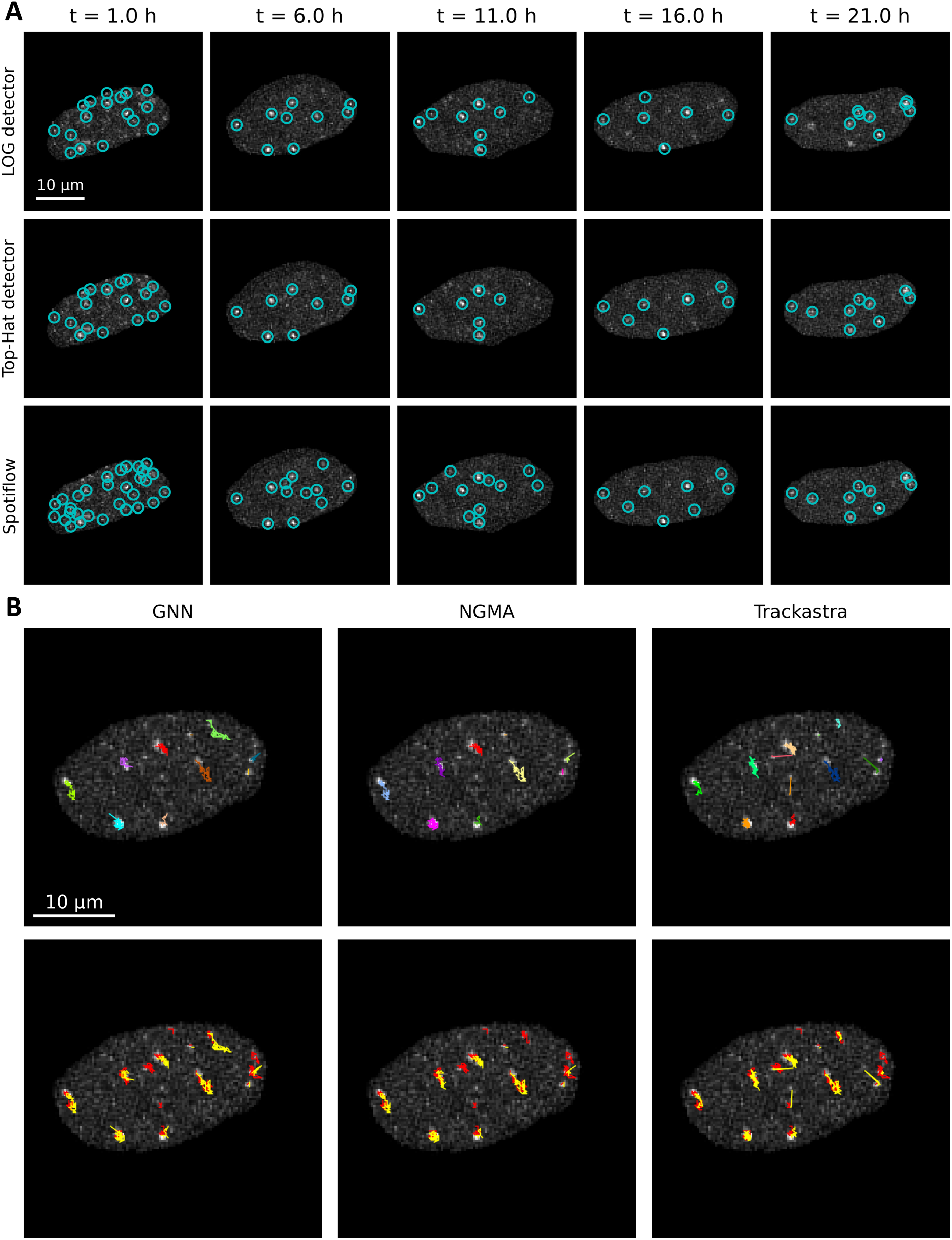
Visual representation of the detected foci and tracks for different algorithms (step 4 and 5). (**A**) Representative images of detected foci using the three tested algorithms at different timepoints. (**B**) The resulting tracks based on the detections of the Top-Hat detector. The top panel indicates the tracks. The bottom panel shows the comparisons of the tracking result (yellow) to the ground-truth tracks (red). In the bottom panel, the ground-truth and predicted tracks were displayed with a small spatial offset to prevent visual overlap. For displaying purposes, the contrast of the images was adjusted.

### Tracking of foci

The final stage of the pipeline is tracking of the foci, for which LiFT includes three alternative algorithms. We implemented a GNN tracker, the NGMA tracker^34^ and the deep-learning tracker Trackastra^35^. As for the nucleus tracking, Trackastra is used as the deep-learning tracker, but with a different pre-trained model. Benchmarking against a manual ground-truth (Figure 5B), showed the best JSC for the tracks was achieved using the Top-Hat detector with NGMA tracking. In contrast, the GNN tracker paired with the Top-Hat detector achieved the best *α* and *β* scores (Table 4). Applying NGMA tracking to the detections of Spotiflow reduced the JSC from 0.533 to 0.449, and a further decrease to 0.390 was observed when the LoG detector was used.

### LiFT applied to irradiated PLC/PRF/5 and U2OS-SSTR2 cells

Using LiFT, we analysed the live-cell imaging data obtained from the PLC/PRF/5 and U2OS-SSTR2 cells following treatment with X-ray irradiation (Table 5). For both cell types, nuclei were segmented with Cellpose-SAM, after which they were tracked with the NND algorithm, applying a minimum track length of 36 frames (3 hours). Subsequently, StackReg was used for the registration step without a preprocessing function. We selected the Top-Hat detector in combination with the GNN tracker, as this resulted in best performance for the PLC/PRF/5 cells (Table 4). The U2OS-SSTR2 cells showed a substantially brighter 53BP1-mClover signal compared to PLC/PRF/5 cells. Consequently, the threshold for the intensity of the local maxima was increased for this cell line, based on representative images, to ensure robust detection performance. Following irradiation, the number of active tracks per nucleus, closely corresponding to the number of detected foci, increased sharply before declining over time as repair progressed (Figure 6A-D, Supplementary Figure 3). In contrast, non-irradiated control samples exhibited a stable number of active tracks throughout the 24 hour observation period for both cell types. Additionally, we investigated track duration (Figure 6E-H). X-ray irradiation increased the dwell time in both cell types, and this increase was significant (*p* < 0.001) relative to the corresponding control (Table 5). No difference in dwell time was detected between the two irradiation doses in PLC/PRF/5 cells (mean difference: 0.4 min; 95% bootstrap CI -2.9 to 3.8; p=0.805). The dwell-time distributions were dominated by short-lived tracks, with a maximum around 30 minutes for both cell types, with a long tail corresponding to a subset of foci that persisted for several hours. As the distribution is predominantly made up from short tracks, the average (and median) dwell-time largely depends on the minimum track length (Supplementary Figure 4). Consequently, this effect should be taken into consideration when tuning the parameters for the tracking of the foci. To assess the impact of track filtering on the derived dwell-time distributions, we excluded tracks of foci already visible at the first frame of a nucleus or persisting until the last frame. Such foci lack well-defined initiation or termination points, and their inclusion may bias dwell-time estimates. This filtering ensured only repair events with clearly defined start and end times contributed to the final dwell-time distributions. For both cell types, the average dwell time decreased slightly across all experimental conditions as a result of excluding these partial tracks (Figure 6I, J).

**Table 5.** Average number of active tracks per cell, the average dwell time, the total number of tracks, and the mean difference in dwell time compared to control (with 95% bootstrap confidence intervals) for the different treatment conditions and cell types. All differences relative to control are significant (*p* < 0.001).

| Cell type | Dose [Gy] | Avg. active tracks per cell | Avg. dwell time [min] | $\Delta$ dwell time to 0 Gy [min] (95% CI) | Number of tracks |
| --- | --- | --- | --- | --- | --- |
| PLC/PRF/5 | 0 | 3.19 | 80.3 | - | 6979 |
|  | 1.5 | 8.80 | 110.9 | 30.6 (26.9–34.3) | 13221 |
|  | 2.5 | 12.74 | 110.5 | 30.2 (26.8–33.6) | 20266 |
| U2OS-SSTR2 | 0 | 7.07 | 75.9 | - | 34568 |
|  | 2.0 | 12.18 | 90.8 | 14.9 (13.4–16.5) | 38987 |

**Figure 6.**
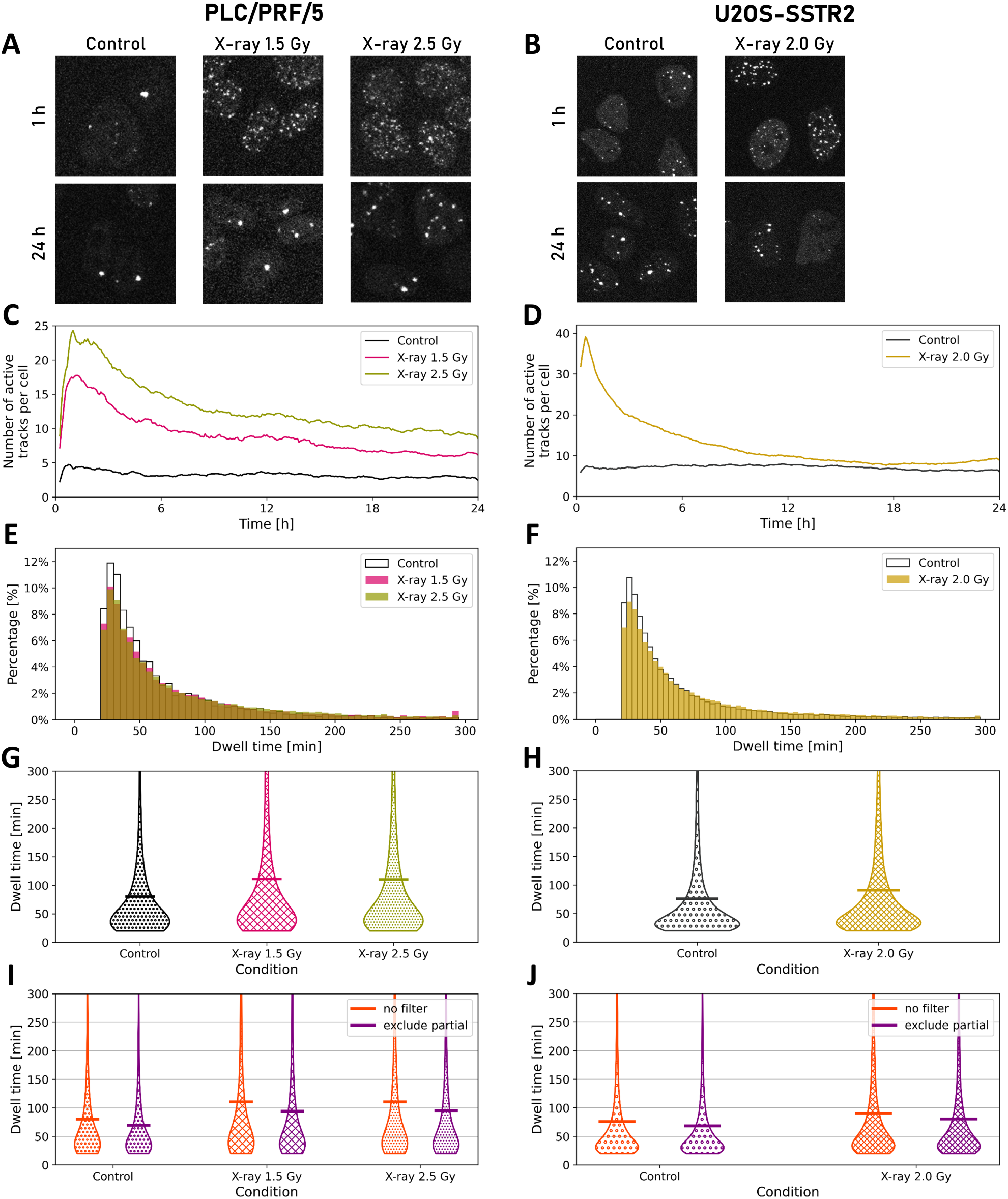
Foci quantification for PLC/PRF/5 and U2OS-SSTR2 cells. The left panels (**A, C, E, G, I**) show results for PLC/PRF/5 cells, and the right panels (**B, D, F, H, J**) for U2OS-SSTR2 cells. (**A**,**B**) Representative images of cells for the different treatment conditions at 1 and 24 hours post X-ray irradiation. For displaying purposes, the contrast of the images was adjusted. (**C**,**D**) Number of active tracks per cell over time. (**E**,**F**) The distribution of obtained track durations (dwell times) for the different treatment conditions. (**G**,**H**) Same data as (**E**,**F**) respectively but represented as a violin plot. (**I, J**) Distribution of track durations when no tracks have been filtered (orange) or when tracks with occluded start or end points have been removed (purple).

## Discussion

In this work, we presented LiFT, an automated and modular software pipeline for the detection, tracking and analysis of DSB foci in live-cell microscopy data. The pipeline resolves the limitations of existing analysis approaches: manual quantification is tedious, time-consuming and subject to operator bias, while the few available automated tools are often closed-source, focusing only on single aspects of the described workflow, or do not leverage recent advances in deep learning. By integrating a wide range of top-performing and state-of-the-art algorithms at every stage, LiFT provides a flexible framework that can be adapted to a wide range of experimental conditions.

A core design principle of LiFT is its modular architecture: because each processing step was implemented as an interchangeable component, individual parts of the pipeline (e.g. the detection and tracking of foci) remain applicable to a broader range of datasets and experimental designs beyond those considered here. For example, the data analysed here were acquired with sampling intervals of 5 or 4 minutes for the PLC/PRF/5 and U2OS-SSTR2 cells, respectively. At substantially higher frame rates, or in cases where cells and foci move more slowly, the registration step included in LiFT may become less relevant and could be omitted without compromising downstream analyses.

Using identical nuclear segmentation masks from step 1, three tracking algorithms in step 2 yielded highly comparable performance. This is most likely attributed to the characteristics of our data, which was acquired with a high temporal resolution (for the cell tracking purposes). Consequently, the inter-frame displacements of the nuclei were minimal, reducing the complexity of the tracking task and limiting the potential for algorithm-specific differences to emerge. Additionally, we did not attempt to resolve “mother–daughter” relationships during cell divisions, as the absence of nuclear stain compromised the reliability of lineage assignment.

Following nucleus tracking, individual cells were cropped from the FOV using their segmentation masks, yielding single-nucleus image stacks followed over time without signal from neighbouring cells. Pixels outside the nuclear mask were set to zero, producing a sharp intensity transition at the nuclear boundary. This masking step ensured accurate registration by eliminating interference from adjacent nuclei. However, we anticipated that the sharp intensity step at the mask border could introduce artefacts during foci detection. In practice, none of the eight detectors implemented in LiFT exhibited this behaviour. While users should remain aware of this potential artefact when extending LiFT with custom detectors, the current implementation remains unaffected.

After isolating the nuclei and performing registration, foci were detected and tracked. Contrary to our expectations based on the particle tracking challenge, the GNN tracker outperformed the NGMA tracker on our dataset^49^. We attribute this to differences in how each algorithm handles the maximum linking distance in combination with gap-closing. Our GNN implementation keeps the maximum linking distance fixed, whereas the NGMA implementation increases it when gap-closing is applied. Because foci undergo largely random diffusion with limited motion and the registration step compensates for nuclear motion, inter-frame displacements in our data are small, and expanding the linking distance during gap-closing can introduce spurious linkages.

Throughout all the steps, LiFT incorporates four deep learning methods (Cellpose-V3, Cellpose-SAM, Spotiflow and Trackastra), each trained on large and diverse datasets to ensure generalizability to new data. Although all of these methods support fine-tuning on user-provided annotations, we deliberately chose not to retrain them within LiFT for two reasons. First, manual annotation is a time-consuming process that would substantially hinder the applicability of the pipeline. Second, fine-tuning on a specific dataset inherently trades generalizability for dataset-specific performance, which risks degrading performance on data from other experimental setups. This trade-off likely explains why Spotiflow did not outperform the traditional Top-Hat detector in our benchmarks: retraining Spotiflow on manual annotations would most likely improve its performance on our specific data, but at the cost of broader applicability. Since LiFT is intended to work across a wide variety of cell lines, microscope types, and experimental conditions, we prioritized the generalizability of the pre-trained models over dataset-specific optimization.

Although the number of foci per nucleus is more commonly reported in the DNA damage literature, we propose using the number of active tracks as a measure of total number of DSBs present at a specific moment. Active tracks comprise foci that persist across multiple frames, making this measure inherently less sensitive to false-positive detections, which typically fail to link consistently over time. Conversely, missed detections in individual frames can still contribute to this metric when gap-closing is enabled, thereby reducing the impact of false negatives. The observed number of active tracks per cell closely follows the established temporal behaviour for the number of foci per nucleus, peaking at approximately 30 minutes post-irradiation after which it decreases^14,51^. Consistent with prior observations, we also detected the expected dose-dependent increase in PLC/PRF/5 cells, with higher irradiation doses yielding larger numbers of active tracks. Together, these results suggest that the number of active tracks is a robust alternative to foci counts, preserving the expected biological trends while being less susceptible to detection artefacts.

Having validated the performance of each step in the pipeline, we validated the measured DSB repair kinetics using a comparison to the literature. The shape of our dwell-time distributions, a prominent peak at short durations with a long tail of persistent events, has also been reported previously^12^. Although average dwell times differed between PLC/PRF/5 and U2OS-SSTR2 cells, both fell within the expected literature range and align with known variability in the DDR across cell lines^52^. For both cell types, dwell times were shorter in non-irradiated controls than after X-ray exposure. This can be explained by the fact that endogenous damage in non-irradiated cells, such as replication-associated lesions, is generally more isolated and less complex than radiation–induced DSBs^53,54^. Accordingly, the increased dwell times observed in irradiated cells reflect this complexity, resulting in longer processing times by the DDR and prolonged track durations. Average track durations were comparable following 1.5 and 2.5 Gy X-ray irradiation in PLC/PRF/5 cells, consistent with prior reports that DSB repair kinetics are largely independent of X-ray dose under similar conditions^55–57^. This is further supported by the temporal evolution of foci numbers, which declined at similar rates for both doses, in agreement with previous observations^57,58^. Previous studies modelled DSB repair times based on *γ*H2AX foci counts, estimating an average repair time of 2.5 hours (range 0.5–4.5 hours)^59^. Bi-exponential models, which partition repair into a fast and a slow component, reported fast-component repair times of 1.18–2.33 hours depending on the beam particle type, with X-rays yielding 1.54 hours^60^. These values are consistent with the dwell times we observed following X-ray irradiation.

Beyond the validation based on previous literature, LiFT offers two methodological advantages over existing analyses. First, our approach measures a full distribution of repair times rather than collapsing the data into two discrete components (fast and slow), which may better capture the underlying biological complexity of the DDR. Second, direct tracking of individual foci extends the applicability of repair-kinetics analysis to treatment modalities where exponential-decay models are not valid. Such models implicitly assume a single, well-defined damage induction event followed by repair, as is the case for acute X-ray exposure. However, treatments such as targeted radionuclide therapy or chemotherapy induce DNA damage continuously over extended periods, causing induction and repair to occur simultaneously. Under these conditions, bulk foci counts cannot be decomposed into induction and repair components, whereas tracking individual foci directly measures the lifetime of each repair event regardless of when it was initiated, providing access to repair kinetics that would otherwise be inaccessible.

A further design consideration of LiFT is its accessibility. The development of computational pipelines for image analysis typically requires considerable programming expertise, limiting their adoption by experimental biologists who would benefit most from them. To address this, we provide LiFT as an open-source Python package alongside a browser-based graphical user interface that exposes the full pipeline without requiring any programming. For users with more technical backgrounds, the same functionality is available through a Jupyter notebook that supports hyperparameter tuning and a command-line interface, supporting integration into automated workflows. By covering this range of interfaces, we aim to make automated foci-tracking analysis accessible to a broad community of researchers, regardless of their computational background.

In conclusion, LiFT enables fully automated, high-accuracy detection and tracking of foci from live-cell imaging data.

By removing the need for manual tracking, which restricted previous studies to relatively small numbers of tracks^13,23^, LiFT allows large foci numbers to be analysed efficiently across diverse experimental conditions. The proposed automation further eliminates the subjective decisions inherent to manual selection, reducing operator bias in determining which foci are included in the analysis. Together, these advances enable LiFT to extract all detectable tracks from time-lapse data, yielding greater statistical power and a more comprehensive characterization of DNA damage dynamics. To make this capability broadly accessible, LiFT is distributed as a freely available open-source Python package with an accompanying graphical user interface, enabling researchers without a background in image analysis or programming to apply the full pipeline to their own data. By providing this generalizable and modular framework, LiFT empowers researchers to investigate the DDR in greater detail and at larger scale, accelerating our understanding of the DNA repair machinery and supporting the development of more effective cancer therapies.

## Acknowledgements

This work was financially supported by the ERC RADIOBIO project [101042537]. We would like to acknowledge Stefan van Alen who performed the imaging of the U2OS-SSTR2 cells. We would like to thank the Erasmus Optical Imaging Centre (OIC) for their assistance and support with the confocal microscope.

## Author contributions statement

T.H.W. conceptualized the study, implemented the software, wrote the manuscript and optimized the microscopy setup. P.E. created and validated the U2OS-SSTR2 cell line and optimized the microscopy setup. J.P. created and validated the the PLC/PRF/5 cells and performed the imaging on these cells. K.H.S. set up the distribution of LiFT, including the Docker container and the Python package. S.F.B.B. supervised the project. I.S. supervised the project, conceptualized the study, and edited the manuscript. J.N. supervised the project, conceptualized the study, edited the manuscript and acquired funding. All authors reviewed the manuscript.

## Additional information

**Accession codes** The code for LiFT is made available on: https://github.com/UU-cellbiology/LiveFoci.

**Competing interests** The authors declare to have no competing interests relevant for this study.

## Notes

### Competing Interest Statement

The authors have declared no competing interest.

## References

1. Barton, M. B. et al. Estimating the demand for radiotherapy from the evidence: A review of changes from 2003 to 2012. Radiother. Oncol. 112, 140–144, DOI: 10.1016/j.radonc.2014.03.024 (2014).

2. van de Kamp, G., Heemskerk, T., Kanaar, R. & Essers, J. DNA double strand break repair pathways in response to different types of ionizing radiation. Front. Genet. Volume 12 - 2021, DOI: 10.3389/fgene.2021.738230 (2021).

3. Harper, J. W. & Elledge, S. J. The DNA damage response: ten years after. Mol. cell 28, 739–745, DOI: 10.1016/j.molcel.2007.11.015 (2007).

4. Deckbar, D., Jeggo, P. A. & Löbrich, M. Understanding the limitations of radiation-induced cell cycle checkpoints. Critical reviews biochemistry molecular biology 46, 271–283, DOI: 10.3109/10409238.2011.575764 (2011).

5. Panier, S. & Boulton, S. J. Double-strand break repair: 53BP1 comes into focus. Nat. reviews Mol. cell biology 15, 7–18, DOI: 10.1038/nrm3719 (2014).

6. Paul, M. W. et al. Role of BRCA2 DNA-binding and C-terminal domain in its mobility and conformation in DNA repair. eLife 10, e67926, DOI: 10.7554/eLife.67926 (2021).

7. Jakl, L., Marková, E., Koláriková, L. & Belyaev, I. Biodosimetry of low dose ionizing radiation using DNA repair foci in human lymphocytes. Genes 11, DOI: 10.3390/genes11010058 (2020).

8. Viau, M. et al. Global quantification of γH2AX as a triage tool for the rapid estimation of received dose in the event of accidental radiation exposure. Mutat. Res. Toxicol. Environ. Mutagen. 793, 123–131, DOI: 10.1016/j.mrgentox.2015.05.009 (2015). Insights into formation and consequences of chromosome aberrations: Report on the 11th International Symposium on Chromosomal Aberrations (ISCA 11), Rhodes, Greece, September 12-14, 2014.

9. Löbrich, M. et al. γH2AX foci analysis for monitoring DNA double-strand break repair: Strengths, limitations and optimization. Cell Cycle 9, 662–669, DOI: 10.4161/cc.9.4.10764 (2010). PMID: 20139725, 10.4161/cc.9.4.10764.

10. Vicar, T. et al. Deepfoci: Deep learning-based algorithm for fast automatic analysis of DNA double-strand break ionizing radiation-induced foci. Comput. Struct. Biotechnol. J. 19, 6465–6480, DOI: 10.1016/j.csbj.2021.11.019 (2021).

11. Heemskerk, T. et al. A novel live-cell microscopy platform for real-time visualization of 53BP1 foci dynamics and accurate dosimetry in proton therapy. Phys. Medica 135, 105020, DOI: 10.1016/j.ejmp.2025.105020 (2025).

12. Roobol, S. J. et al. Comparison of high- and low-LET radiation-induced DNA double-strand break processing in living cells. Int. J. Mol. Sci. 21, DOI: 10.3390/ijms21186602 (2020).

13. Jakob, B., Splinter, J., Durante, M. & Taucher-Scholz, G. Live cell microscopy analysis of radiation-induced DNA double-strand break motion. Proc. Natl. Acad. Sci. 106, 3172–3177, DOI: 10.1073/pnas.0810987106 (2009). https://www.pnas.org/doi/pdf/10.1073/pnas.0810987106.

14. Sollazzo, A. et al. Live dynamics of 53BP1 foci following simultaneous induction of clustered and dispersed DNA damage in U2OS cells. Int. J. Mol. Sci. 19, DOI: 10.3390/ijms19020519 (2018).

15. Boudreau, C. et al. Excitation light dose engineering to reduce photo-bleaching and photo-toxicity. Sci. reports 6, 30892, DOI: 10.1038/srep30892 (2016).

16. Zhang, Y. et al. A Poisson-Gaussian denoising dataset with real fluorescence microscopy images. In 2019 IEEE/CVF Conference on Computer Vision and Pattern Recognition (CVPR), 11702–11710, DOI: 10.1109/CVPR.2019.01198 (2019).

17. Laine, R. F., Jacquemet, G. & Krull, A. Imaging in focus: an introduction to denoising bioimages in the era of deep learning. The international J. biochem. – cell biology 140, 106077, DOI: 10.1016/j.biocel.2021.106077 (2021).

18. Schirripa Spagnolo, C. & Luin, S. Impact of temporal resolution in single particle tracking analysis. Discov. Nano 19, 87, DOI: 10.1186/s11671-024-04029-1 (2024).

19. Jaqaman, K. & Danuser, G. Computational image analysis of cellular dynamics: a case study based on particle tracking. Cold Spring Harb. Protoc. 2009, pdb–top65, DOI: 10.1101/pdb.top65 (2009).

20. Smal, I., Loog, M., Niessen, W. & Meijering, E. Quantitative comparison of spot detection methods in fluorescence microscopy. IEEE Transactions on Med. Imaging 29, 282–301, DOI: 10.1109/TMI.2009.2025127 (2010).

21. Ivashkevich, A., Redon, C. E., Nakamura, A. J., Martin, R. F. & Martin, O. A. Use of the γ-H2AX assay to monitor DNA damage and repair in translational cancer research. Cancer Lett. 327, 123–133, DOI: 10.1016/j.canlet.2011.12.025 (2012). Special Issue: Oxidative Stress-Based Cancer Biomarkers.

22. Rothkamm, K. et al. Laboratory intercomparison on the γ-H2AX foci assay. Radiat. research 180, 149–155, DOI: 10.1667/RR3238.1 (2013).

23. Dolan, D., Nelson, G., Zupanic, A., Smith, G. & Shanley, D. Systems modelling of NHEJ reveals the importance of redox regulation of ku70/80 in the dynamics of DNA damage foci. PloS one 8, e55190, DOI: 10.1371/journal.pone.0055190 (2013).

24. Segeren, H. A., van Liere, E. A., Riemers, F. M., de Bruin, A. & Westendorp, B. Oncogenic RAS sensitizes cells to drug-induced replication stress via transcriptional silencing of P53. Oncogene 41, 2719–2733, DOI: 10.1038/s41388-022-02291-0 (2022).

25. Georgescu, W. et al. Characterizing the DNA damage response by cell tracking algorithms and cell features classification using high-content time-lapse analysis. PLoS One 10, e0129438, DOI: 10.1371/journal.pone.0129438 (2015).

26. Bucevičius, J., Lukinavičius, G. & Gerasimaitė, R. The use of Hoechst dyes for DNA staining and beyond. Chemosensors 6, DOI: 10.3390/chemosensors6020018 (2018).

27. Sen, O., Saurin, A. T. & Higgins, J. M. The live cell DNA stain SiR-Hoechst induces DNA damage responses and impairs cell cycle progression. Sci. reports 8, 7898, DOI: 10.1038/s41598-018-26307-6 (2018).

28. Perrin, J. et al. Radiobiological comparison of holmium-166 and yttrium-90 for hepatocellular carcinoma treatment in vitro. Nucl. Medicine Commun. DOI: 10.1097/MNM.0000000000002190 (2026).

29. Stringer, C., Wang, T., Michaelos, M. & Pachitariu, M. Cellpose: a generalist algorithm for cellular segmentation. Nat. methods 18, 100–106, DOI: 10.1038/s41592-020-01018-x (2021).

30. Pachitariu, M., Rariden, M. & Stringer, C. Cellpose-SAM: superhuman generalization for cellular segmentation. bioRxiv DOI: 10.1101/2025.04.28.651001 (2025). https://www.biorxiv.org/content/early/2025/05/01/2025.04.28.651001.full.pdf.

31. Kirillov, A. et al. Segment anything. In 2023 IEEE/CVF International Conference on Computer Vision (ICCV), 3992–4003, DOI: 10.1109/ICCV51070.2023.00371 (2023).

32. Starck, J.-L., Elad, M. & Donoho, D. Redundant multiscale transforms and their application for morphological component separation. Adv. Imaging Electron Phys. 287–348, DOI: 10.1016/S1076-5670(04)32006-9 (2004).

33. Mallat, S. A wavelet tour of signal processing (Elsevier, 1999).

34. Reuter, M. et al. BRCA2 diffuses as oligomeric clusters with RAD51 and changes mobility after DNA damage in live cells. J. Cell Biol. 207, 599–613, DOI: 10.1083/jcb.201405014 (2014). https://rupress.org/jcb/article-pdf/207/5/599/1586090/jcb_201405014.pdf.

35. Gallusser, B. & Weigert, M. Trackastra: Transformer-based cell tracking for live-cell microscopy. In European conference on computer vision, 467–484, DOI: 10.1007/978-3-031-73116-7_27 (Springer, 2024).

36. Maška, M. et al. The cell tracking challenge: 10 years of objective benchmarking. Nat. Methods 20, 1010–1020, DOI: 10.1038/s41592-023-01879-y (2023).

37. Thevenaz, P., Ruttimann, U. & Unser, M. A pyramid approach to subpixel registration based on intensity. IEEE Transactions on Image Process. 7, 27–41, DOI: 10.1109/83.650848 (1998).

38. Klein, S., Staring, M., Murphy, K., Viergever, M. A. & Pluim, J. P. W. elastix: A toolbox for intensity-based medical image registration. IEEE Transactions on Med. Imaging 29, 196–205, DOI: 10.1109/TMI.2009.2035616 (2010).

39. Bakshi, A., Indyk, P., Jayaram, R., Silwal, S. & Waingarten, E. Near-linear time algorithm for the chamfer distance. Adv. Neural Inf. Process. Syst. 36, 66833–66844 (2023).

40. Figueiredo, M. A. & Nowak, R. D. Wavelet-based image estimation: an empirical Bayes approach using Jeffrey’s noninformative prior. Trans. Img. Proc. 10, 1322–1331, DOI: 10.1109/83.941856 (2001).

41. Sage, D., Neumann, F., Hediger, F., Gasser, S. & Unser, M. Automatic tracking of individual fluorescence particles: application to the study of chromosome dynamics. IEEE Transactions on Image Process. 14, 1372–1383, DOI: 10.1109/TIP.2005.852787 (2005).

42. Lengert, N. et al. AutoFoci, an automated high-throughput foci detection approach for analyzing low-dose DNA double-strand break repair. Sci. Reports 8, 17282, DOI: 10.1038/s41598-018-35660-5 (2018).

43. Lapytsko, A., Kollarovic, G., Ivanova, L., Studencka, M. & Schaber, J. FoCo: a simple and robust quantification algorithm of nuclear foci. BMC bioinformatics 16, 392, DOI: 10.1186/s12859-015-0816-5 (2015).

44. Vincent, L. Morphological grayscale reconstruction in image analysis: applications and efficient algorithms. IEEE Transactions on Image Process. 2, 176–201, DOI: 10.1109/83.217222 (1993).

45. Smal, I., Niessen, W. & Meijering, E. A new detection scheme for multiple object tracking in fluorescence microscopy by joint probabilistic data association filtering. In 2008 5th IEEE International Symposium on Biomedical Imaging: From Nano to Macro, 264–267, DOI: 10.1109/ISBI.2008.4540983 (2008).

46. Rezatofighi, S. H., Hartley, R. & Hughes, W. E. A new approach for spot detection in total internal reflection fluorescence microscopy. In 2012 9th IEEE International Symposium on Biomedical Imaging (ISBI), 860–863, DOI: 10.1109/ISBI.2012.6235684 (2012).

47. Dominguez Mantes, A. et al. Spotiflow: accurate and efficient spot detection for fluorescence microscopy with deep stereographic flow regression. Nat. methods 22, 1495–1504, DOI: 10.1038/s41592-025-02662-x (2025).

48. Jaqaman, K. et al. Robust single-particle tracking in live-cell time-lapse sequences. Nat. methods 5, 695–702, DOI: 10.1038/nmeth.1237 (2008).

49. Chenouard, N. et al. Objective comparison of particle tracking methods. Nat. methods 11, 281–289, DOI: 10.1038/nmeth.2808 (2014).

50. Ershov, D. et al. Trackmate 7: integrating state-of-the-art segmentation algorithms into tracking pipelines. Nat. methods 19, 829–832, DOI: 10.1038/s41592-022-01507-1 (2022).

51. Schultz, L. B., Chehab, N. H., Malikzay, A. & Halazonetis, T. D. P53 binding protein 1 (53BP1) is an early participant in the cellular response to DNA double-strand breaks. J. Cell Biol. 151, 1381–1390, DOI: 10.1083/jcb.151.7.1381 (2000). https://rupress.org/jcb/article-pdf/151/7/1381/1863179/0008067.pdf.

52. Firsanov, D., Vasilishina, A., Kropotov, A. & Mikhailov, V. Dynamics of γH2AX formation and elimination in mammalian cells after X-irradiation. Biochimie 94, 2416–2422, DOI: 10.1016/j.biochi.2012.06.019 (2012). Special Section on Epigenetics.

53. Jakob, B. et al. Differential repair protein recruitment at sites of clustered and isolated DNA double-strand breaks produced by high-energy heavy ions. Sci. reports 10, 1443, DOI: 10.1038/s41598-020-58084-6 (2020).

54. Nickoloff, J. A., Sharma, N. & Taylor, L. Clustered DNA double-strand breaks: Biological effects and relevance to cancer radiotherapy. Genes 11, DOI: 10.3390/genes11010099 (2020).

55. Lee, U.-S.Lee, D.-H. & Kim, E.-H. Characterization of γ-H2AX foci formation under alpha particle and X-ray exposures for dose estimation. Sci. reports 12, 3761, DOI: 10.1038/s41598-022-07653-y (2022).

56. Asaithamby, A. & Chen, D. J. Cellular responses to DNA double-strand breaks after low-dose γ-irradiation. Nucleic Acids Res. 37, 3912–3923, DOI: 10.1093/nar/gkp237 (2009). https://academic.oup.com/nar/article-pdf/37/12/3912/16751904/gkp237.pdf.

57. Horn, S., Barnard, S. & Rothkamm, K. γ-H2AX-based dose estimation for whole and partial body radiation exposure. PloS one 6, e25113, DOI: 10.1371/journal.pone.0025113 (2011).

58. Mariotti, L. G. et al. Use of the γ-H2AX assay to investigate DNA repair dynamics following multiple radiation exposures. PloS one 8, e79541, DOI: 10.1371/journal.pone.0079541 (2013).

59. Scott, B. R. Modeling DNA double-strand break repair kinetics as an epiregulated cell-community-wide (Epicellcom) response to radiation stress. Dose-Response 9, dose–response.10–039.Scott, DOI: 10.2203/dose-response.10-039.Scott (2011). PMID: 22461762, https://doi.org/10.2203/dose-response.10-039.Scott.

60. Plante, I., Slaba, T., Shavers, Z. & Hada, M. A bi-exponential repair algorithm for radiation-induced double-strand breaks: Application to simulation of chromosome aberrations. Genes 10, DOI: 10.3390/genes10110936 (2019).

