## supplemental information for "LiFT: Live foci tracking for quantitative analysis of DNA damage dynamics"

### Supplementary material

#### Cell-lines and data acquisition

DNA double strand breaks (DSBs) were visualized in two cell lines, PLC/PRF/5 and U2OS-SSTR2, each expressing an endogenous fluorescently tagged 53BP1-mClover protein. The PLC/PRF/5_mClover-53BP1 cells were developed and validated previously[1]. Briefly, knock-ins were generated using CRISPR-Cas9 technology, and single-cell clones obtained by limiting dilution were validated by genomic PCR, immunofluorescence staining and clonogenic survival. The U2OS-SSTR2-mClover-53BP1 line was developed for this study using the same procedure. Prior to imaging, DSBs were induced by X-ray irradiation (0, 1.5 or 2.5 Gy for PLC/PRF/5; 0 or 2 Gy for U2OS-SSTR2) using an X-strahl RS320 cabinet irradiator (195 kV, 10 mA, 1.6 Gy/minute). Imaging was performed the following 24 hours in 8-well plates (ibidi) on a Leica TCS SP8 confocal microscope equipped with a cage incubator (Okolab) maintaining 37 °C, 5% $CO_2$ and a humid atmosphere. Imaging was performed with a 40× immersion objective, a 488 nm excitation laser and a Photomultiplier tube (PMT) detector, acquiring z-stacks with a pixel size of 284 × 284 nm (1024 × 1024 pixels) and a z-step of 750 nm. Two biological replicates were acquired for PLC/PRF/5 and three for U2OS-SSTR2. To emulate distinct imaging setups, acquisition parameters were deliberately varied between the two cell lines: PLC/PRF/5 cells were imaged every 5 minutes, U2OS-SSTR2 cells every 4 minutes, and the laser power, detector gain and detector bandwidth were also altered, yielding datasets with distinct image characteristics (Supplementary Table 1). For all analyses, the acquired z-stacks were converted to 2D images using maximum-intensity projection, following the protocol in the study by Perrin *et al*[1].

**Supplementary Table 1.** Image acquisition parameters for the different cell types.

| Cell type | sampling time [min] | laser power [%] | detector gain | detection bandwidth [nm] |
|---|---|---|---|---|
| PLC//PRF/5 | 5 | 0.2 | 898 | 500-550 |
| U2OS-SSTR2 | 4 | 0.3 | 964 | 500-570 |

### References


**1.** Perrin, J. *et al.* Radiobiological comparison of holmium-166 and yttrium-90 for hepatocellular carcinoma treatment in vitro. *Nucl. Medicine Commun.* DOI: https://doi.org/10.1097/MNM.0000000000002190 (2026).

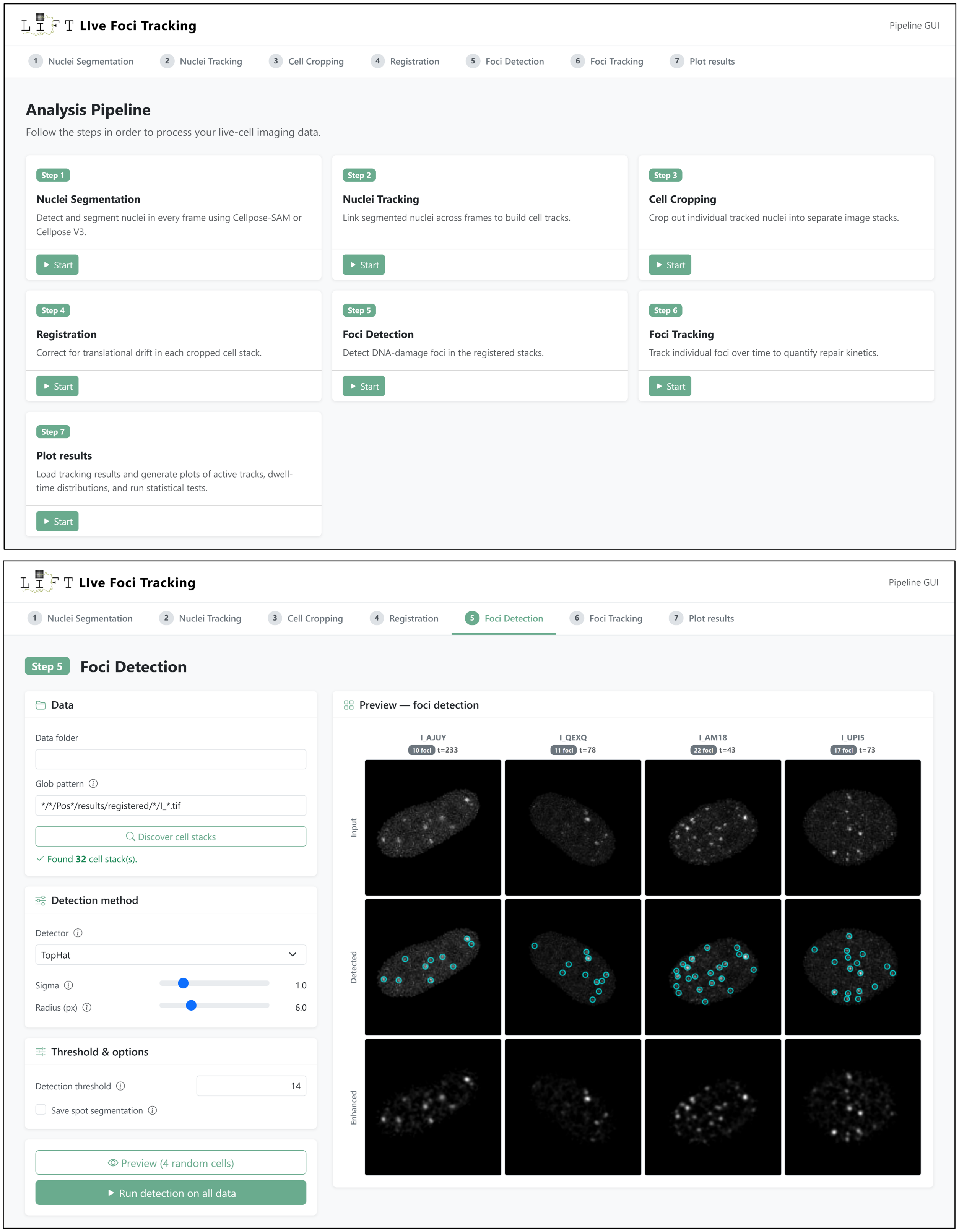


**Supplementary Figure 1.** Screenshots of the interactive browser-based graphical user interface. The top panel depicts the main page with all the steps implemented in LiFT. The bottom panel depict the foci detection stage with the detection methods and associated parameters on the left, and the resulting detections on the right.

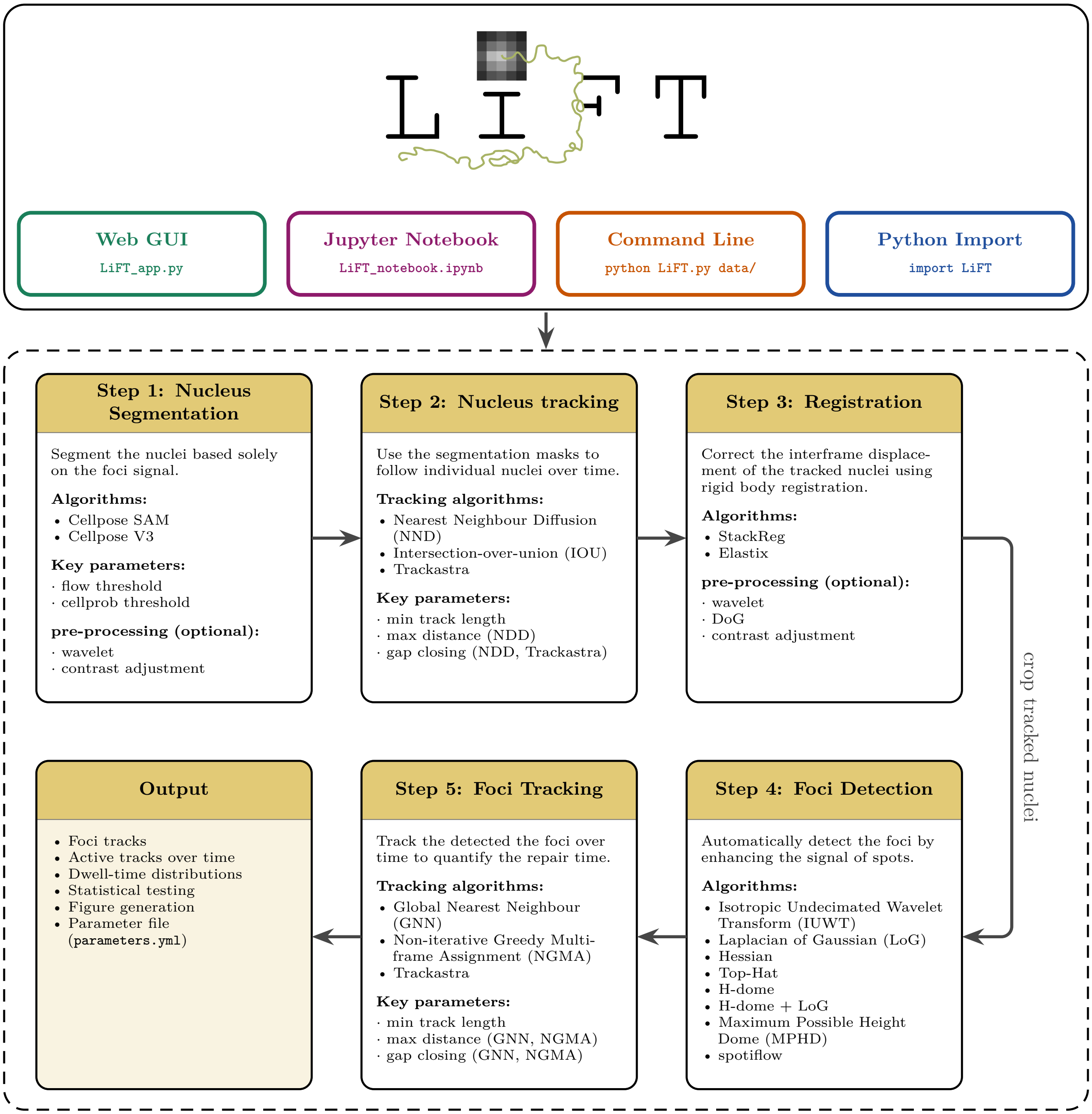


**Supplementary Figure 2.** Schematic overview of the LiFT pipeline and its components. The top panel shows the four ways in which LiFT can be run: a browser-based graphical user interface, a Jupyter notebook, a command-line interface, and as an imported Python module. The bottom panel shows the five processing steps of the pipeline together with the alternative algorithms and key parameters available for each step. The final output is summarized in the bottom left.

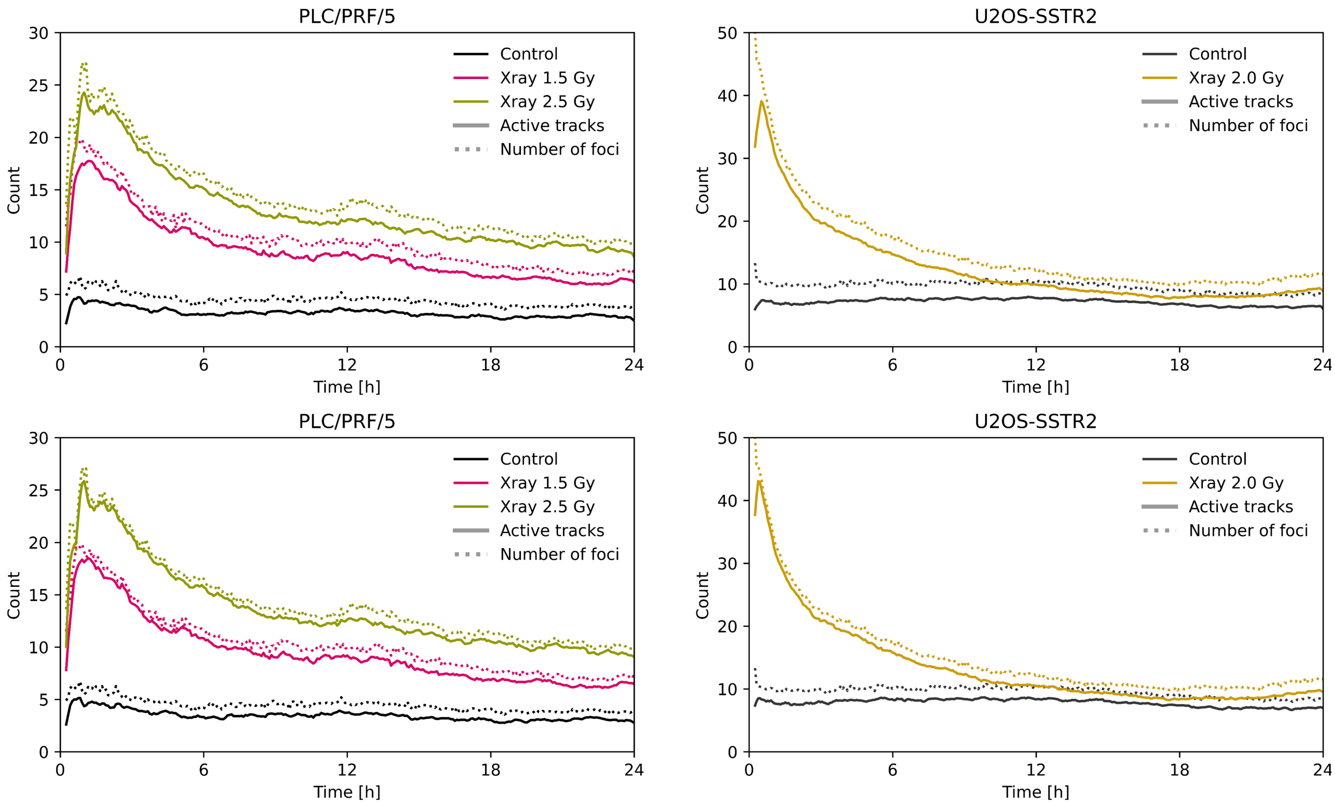


**Supplementary Figure 3.** Comparison of active tracks and detected foci counts. The top panel shows this comparison using a minimum track length of 20 minutes (four detections for PLC/PRF/5 cells and five detections for U2OS-SSTR2 cells). Lowering the minimum track-length progressively incorporates additional short-lived detections into the active-track count, which may include transient or false-positive events. To illustrate this effect, the bottom panel shows the same comparison using a minimum of three detections, in which the difference between the active tracks and detected foci counts decreased.

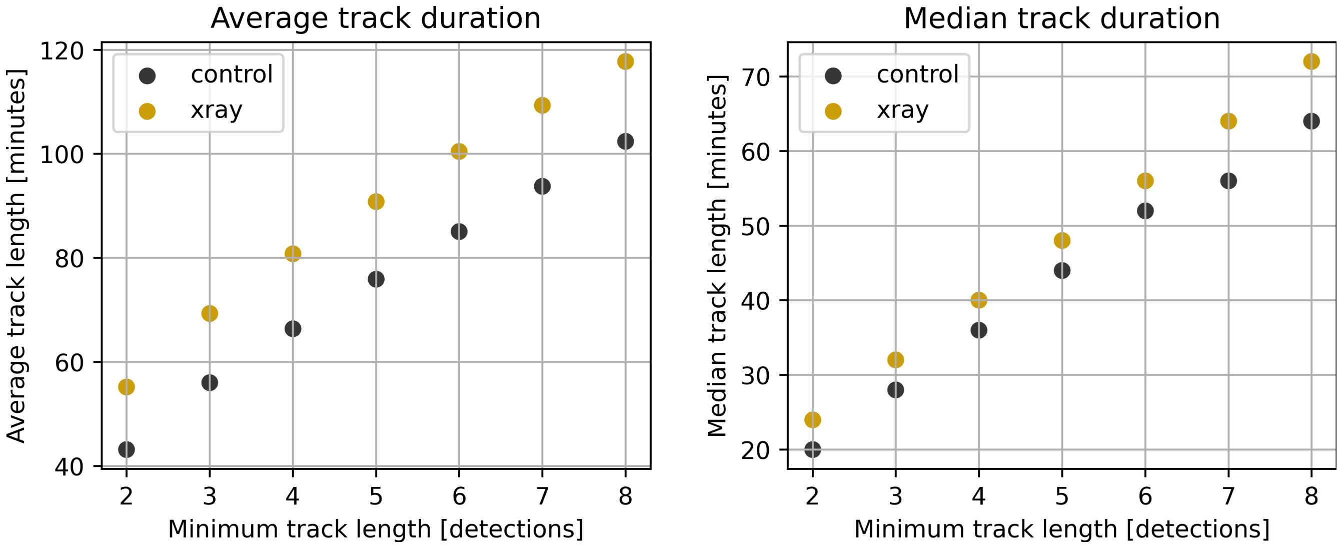


**Supplementary Figure 4.** Dependence of average (left) and median (right) track length on the minimum track-length threshold used for tracking in the U2OS-SSTR2 cells. Increasing the minimum required track length progressively excludes short tracks, leading to an increase in both the measured average and median track lengths.
